# Antagonistic paralogs control a switch between growth and pathogen resistance in *C. elegans*

**DOI:** 10.1101/357756

**Authors:** Kirthi C. Reddy, Tal Dror, Ryan S. Underwood, Guled A. Osman, Christopher A. Desjardins, Christina A. Cuomo, Michalis Barkoulas, Emily R. Troemel

**Affiliations:** Division of Biological Sciences, University of California, San Diego, La Jolla, CA 92093 USA; Infectious Disease and Microbiome Program, Broad Institute, Cambridge MA 02142 USA; Department of Life Sciences, Imperial College, London SW7 2AZ, UK

## Abstract

Immune genes are under intense pressure from pathogens, which cause these genes to diversify over evolutionary time and become species-specific. Through a forward genetic screen we recently described a *C. elegans*-specific gene called *pals-22* to be a repressor of “Intracellular Pathogen Response” or IPR genes. Here we describe *pals-25*, which, like *pals-22*, is a species-specific gene of unknown biochemical function. We identified *pals-25* in a screen for suppression of *pals-22* mutant phenotypes and found that mutations in *pals-25* suppress all known phenotypes caused by mutations in *pals-22*. These phenotypes include increased IPR gene expression, thermotolerance, and immunity against natural pathogens. Mutations in *pals-25* also reverse the reduced lifespan and slowed growth of *pals-22* mutants. Transcriptome analysis indicates that *pals-22* and *pals-25* control expression of genes induced not only by natural pathogens of the intestine, but also by natural pathogens of the epidermis. Indeed, in an independent forward genetic screen we identified *pals-22* as a repressor and *pals-25* as an activator of epidermal defense gene expression. These phenotypic and evolutionary features of *pals-22* and *pals-25* are strikingly similar to species-specific R gene pairs in plants that control immunity against co-evolved pathogens.

## Introduction

Evolutionarily ancient genes control core processes in diverse organisms. For example, the >500 million-year-old Hox gene cluster is required for establishing body plan polarity in animals as diverse as worms, flies and humans [1]. However, evolutionarily young genes can also play key roles in development. For example, the *Drosophila* Umbrea gene only evolved within the *Drosophila* lineage in the last 15 million years but is essential for chromosome segregation in *Drosophila melanogaster* [2]. In general, the functions of evolutionarily young genes are less well understood than the function of evolutionarily ancient genes.

New genes can arise through gene duplication and diversification [3]. Extensive gene duplication can lead to large, expanded gene families, which appear ‘species-specific’ if there is significant diversification away from the ancestral gene. The function of species-specific genes can provide insight into the pressures imposed upon organisms in their recent evolutionary past. Pathogen infection imposes some of the strongest selective pressure on organisms, and accordingly, many species-specific, expanded gene families are involved in immunity. One example is the family of mouse Naip genes, which encode sensor proteins in the inflammasome that detect bacteria to trigger cytokine release and cell death [4]. Another example is the plant R genes, which detect virulence factors from co-evolved pathogens to activate effector-triggered immunity [5]. Interestingly, a growing theme in plant R genes is that they can function as opposing gene pairs, with one R gene promoting host defense and the other R gene inhibiting host defense. Of note, both the Naip and R genes were identified through unbiased forward genetic screens for immune genes.

Recently, we described a forward genetic screen in *C. elegans* for genes that regulate the transcriptional response to natural intracellular pathogens [6]. From this screen we identified a *C. elegans*-specific gene called *pals-22* that regulates expression of Intracellular Pathogen Response or IPR genes. Interestingly, we found that *pals-22* also regulates proteostasis, potentially through ubiquitin ligase activity (see below). The ‘*pals’* signature stands for <u>p</u>rotein containing <u>ALS</u>2CR12 signature, which is found in the single *pals* gene in humans called ALS2CR12. A genome-wide association study implicated ALS2CR12 in amyotrophic lateral sclerosis (ALS) [7], although this gene has no known role in ALS, and its biological function is unknown. The *pals* gene family has only a single member each in the mouse and human genomes, but is substantially expanded in *Caenorhabditis* genomes: *C. elegans* has 39 *pals* genes; *C. remanei* has 18 *pals* genes; *C. brenneri* has 8 *pals* genes; and *C. briggsae* has 8 *pals* genes [8].

*pals-22* mutants have several striking phenotypes in *C. elegans*. First, *pals-22* mutants have constitutive expression of several IPR genes including the cullin gene *cul-6*, which is predicted to encode a component of a Cullin-Ring Ligase complex. Second, *pals-22* mutants have increased tolerance of proteotoxic stressors, and this increased tolerance requires the wild-type function of *cul-6*. Third, *pals-22* mutants have less robust health in the absence of stressors. In particular, they have slowed development and shorter lifespans compared to wild-type animals. Fourth, as shown by another group who identified *pals-22* in an independent forward genetic screen, *pals-22* mutants have increased transgene silencing, and increased RNA interference (RNAi) against exogenous RNA [8]. Thus, loss-of-function mutations in *pals-22* appear to broadly reprogram the physiology of *C. elegans*.

Here we describe a forward genetic screen for suppressors of *pals-22* and identify another *pals* gene called *pals-25*. Interestingly, although it appears that *pals-25* and *pals-22* are in an operon together, these two genes function antagonistically and direct opposing phenotypes. We show that mutations in *pals-25* strongly suppress all the physiological phenotypes seen in *pals-22* mutants, including IPR gene expression, stress resistance, lifespan, development and transgene silencing. Furthermore, we find that *pals-22* mutants have increased resistance against natural intracellular pathogens, like the microsporidian species *Nematocida parisii* and the Orsay virus. This increased resistance is suppressed by mutations in *pals-25*. Also, we use RNA-seq analysis to show that the *pals-22/pals-25* gene pair (hereafter referred to as *pals-22/25*) regulate expression of a majority of the genes induced by natural pathogens of the intestine and find that most of these genes are also induced by blockade of the proteasome. Interestingly, we observe that *pals-22* and *pals-25* also regulate expression of genes induced by natural eukaryotic pathogens infecting through the epidermis. Indeed, in an independent forward genetic screen to find regulators of epidermal defense gene expression we identified additional mutant alleles of *pals-22* and *pals-25*. In summary, the species-specific *pals-22/25* gene pair control an entire physiological program that balances growth with increased proteostasis capacity and resistance against diverse natural pathogens.

## Results

### *pals-25* is required for increased IPR gene expression in *pals-22* mutants

Previously we found that wild-type *pals-22* represses expression of IPR genes: *pals-22* mutants have constitutive expression of several IPR genes including *pals-5* [6]. A transcriptional reporter consisting of the 1273 bp upstream region of *pals-5* fused to GFP, *pals-5p::GFP*, is a reliable marker of IPR gene expression [9] and exhibits strong GFP expression in a *pals-22* mutant background [6] (Fig 1A-C). To find positive regulators of IPR gene expression, we mutagenized *pals-22; pals-5p::GFP* strains and screened for loss of GFP expression in F2 animals. From one screen in the *pals-22(jy1)* mutant background and one screen in the *pals-22(jy3)* mutant background we screened a total of ~23,000 haploid genomes and found eight independent mutant alleles that almost entirely reverse the increased *pals-5p::GFP* expression back to wild-type levels in *pals-22* mutants (Fig 1A-F). All of these alleles are recessive, segregate in Mendelian ratios, and fail to complement each other. These results suggest they all have loss-of-function mutations in the same gene.

**Fig 1.**
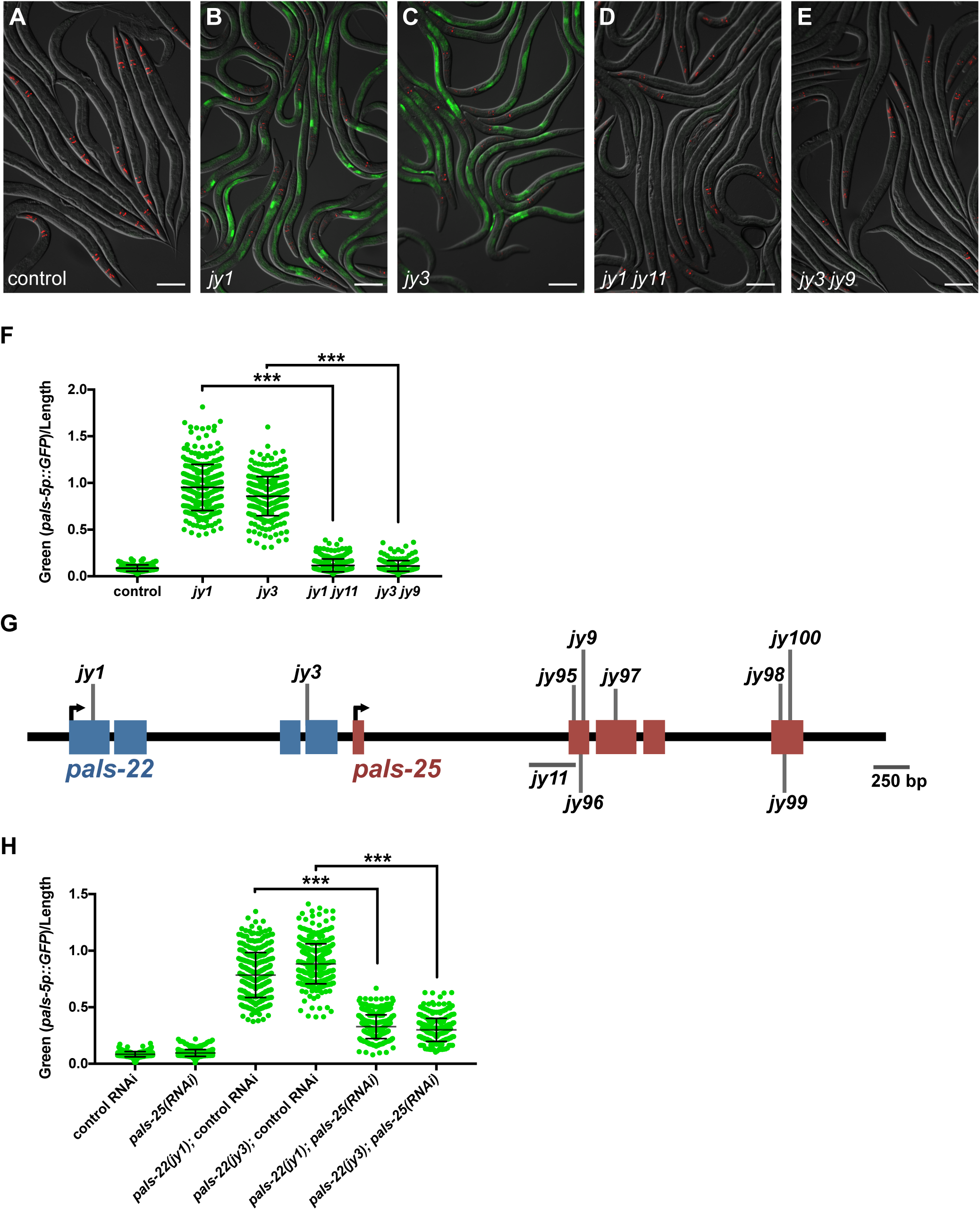
*pals-25* is required for increased *pals-5p::GFP* expression in *pals-22* mutants. (A-E) Mutants isolated from *pals-22* suppressor screens show decreased expression of the *pals-5p::GFP* reporter. Shown are (A) wild-type, (B) *pals-22(jy1)*, (C) *pals-22(jy3)*, (D) *pals-22(jy1) pals-25(jy11)*, and (E) *pals-22(jy3) pals-25(jy9)* animals. Green is *pals-5p::GFP*, red is *myo-2p::mCherry* expression in the pharynx as a marker for presence of the transgene. Images are overlays of green, red and Nomarski channels and were taken with the same camera exposure for all. Scale bar, 100 *µ*m. (F) *pals-5p::GFP* expression quantified in *pals-22* suppressor mutants using a COPAS Biosort machine to measure the mean GFP signal and length of individual animals, indicated by green dots. Mean signal of the population is indicated by black bars, with error bars as SD. Graph is a compilation of three independent replicates, with at least 100 animals analyzed in each replicate. *** p < 0.001 with Student’s t-test. (G) *pals-22* and *pals-25* gene coding structure (UTR not shown), with blue exons for *pals-22* and red exons for *pals-25*. See S1 Table for residues altered. (H) *pals-5p::GFP* expression in animals treated with either L4440 RNAi control or *pals-25* RNAi, quantified using a COPAS Biosort machine to measure the mean GFP signal and length of animals. Parameters the same as in (F).

We used whole-genome sequencing of two mutant strains (*jy9* and *jy100*) to identify the causative alleles [10] and found predicted loss of function mutations in *pals-25* in both strains. Further sequencing identified *pals-25* mutations in the remaining six mutant strains (Fig 1G, S1 Table). *pals-25* appears to be in an operon just downstream of *pals-22*, and while these two genes are paralogs, they share limited sequence similarity, with no significant similarity on the DNA level and only 19.4% identity on the amino acid level. Of note, neither *pals-22* nor *pals-25* have obvious orthologs in other *Caenorhabditis* species, and thus appear to be specific to *C. elegans* [8]. To further confirm that *pals-25* regulates *pals-5p::GFP* gene expression in *pals-22* mutants, we performed RNAi against *pals-25* in a *pals-22; pals-5p::GFP* strain. As expected, we found lowered expression of *pals-5p::GFP* (Fig 1H, S2 Fig), indicating that wild-type *pals-25* is required for the increased expression of *pals-5p::GFP* seen in a *pals-22* mutant background.

These observations suggest that *pals-25* acts downstream of *pals-22* to activate mRNA expression of IPR genes. To test this hypothesis, we used qRT-PCR to measure levels of endogenous mRNA in *pals-22 pals-25* mutants compared to *pals-22* mutants and wild-type animals (Fig 2A). We analyzed mRNA levels of *pals-5*, as well as seven other IPR genes including nematode-specific genes of unknown function (F26F2.1, F26F2.3, and F26F2.4), and predicted ubiquitin ligase components *skr-3, skr-4, skr-5* and *cul-6*. Here we found that mutations in *pals-25* reverse the elevated mRNA levels of all eight of these IPR genes in a *pals-22* mutant background back to near wild-type levels. Importantly, a non-IPR gene, *skr-1*, is not affected by mutations in *pals-22* or *pals-25*. These results indicate that in a *pals-22* mutant background, wild-type *pals-25* activates IPR gene expression.

**Fig 2.**
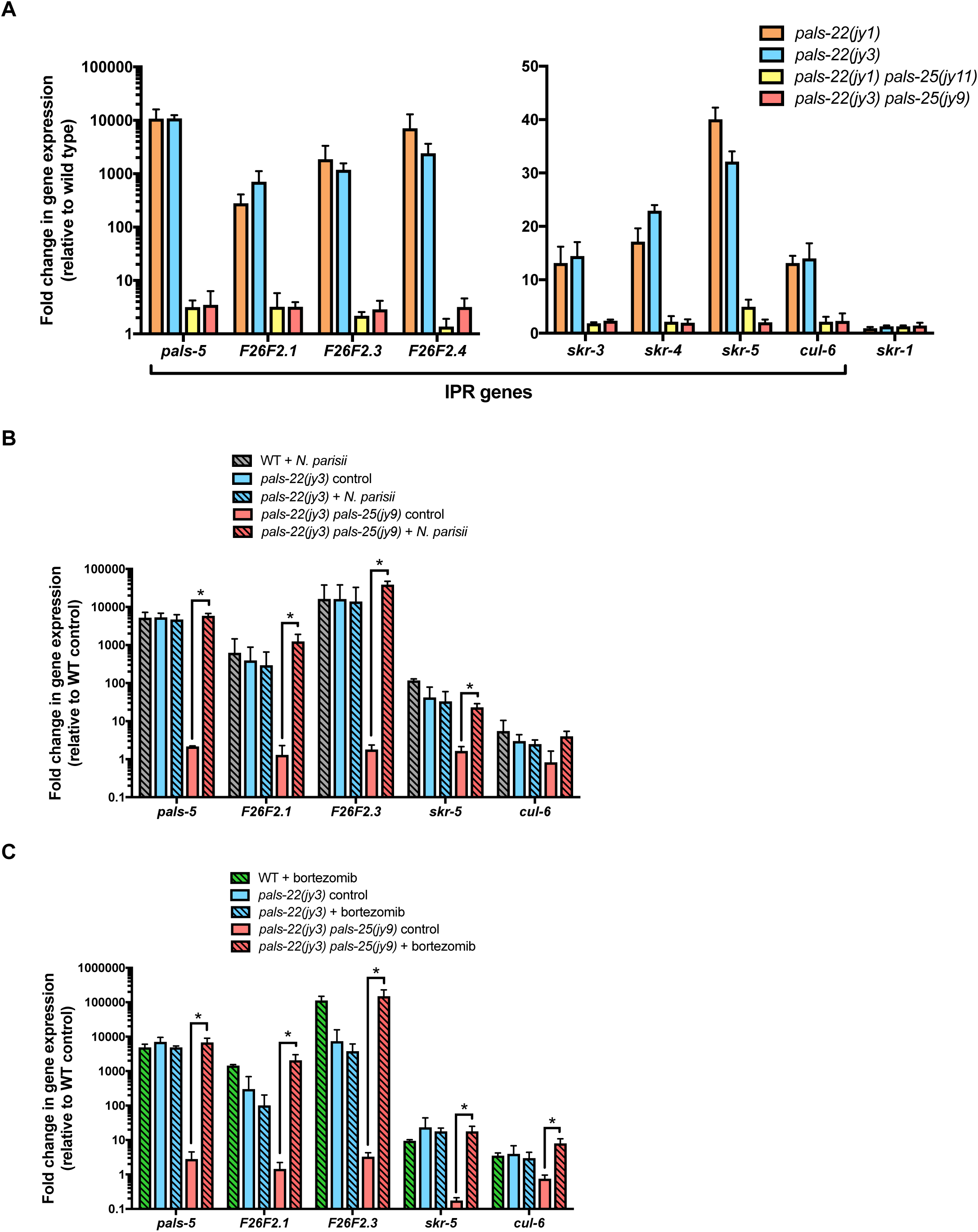
*pals-25* is required for increased IPR gene expression in *pals-22* mutants, but not for IPR induction in response to infection or proteasome inhibition. (A) qRT-PCR measurement of gene expression in *pals-22* and *pals-22 pals-25* animals, shown as the fold change relative to wild-type control. (B-C) qRT-PCR measurement of IPR gene expression in *pals-22(jy3)* and *pals-22(jy3) pals-25(jy9)* animals following 4 hours of infection with *N. parisii* (B) or treatment with the proteasome inhibitor bortezomib (C). For (A-C), results shown are the average of two independent biological replicates and error bars are SD. * p < 0.05 with Student’s t-test.

Previous analysis indicated that *pals-22* is broadly expressed in several tissues in the animal, including the intestine and the epidermis [6, 8]. Similarly, we found that *pals-25* is broadly expressed. Using a fosmid containing *pals-25* with endogenous *cis* regulatory control and tagged at the C terminus with GFP and 3xFLAG [11], we observed PALS-25::GFP expression throughout the animal, including expression in the neurons, epidermis, and intestine (S2B Fig). We did not see any change in PALS-25::GFP expression after *pals-22* RNAi treatment (S2C Fig).

### IPR genes are induced by infection and by proteasome blockade in *pals-22 pals-25* mutants

As *pals-25* is required to activate IPR gene expression in a *pals-22* mutant background, we wondered whether *pals-25* was required for inducing IPR gene expression in response to external triggers. We originally identified IPR genes because of their induction by *N. parisii* infection [6, 9], which is an intracellular pathogen in the Microsporidia phylum that invades and undergoes its entire replicative life cycle inside *C. elegans* intestinal cells [12]. We therefore infected animals with *N. parisii* and compared induction of IPR genes in *pals-22 pals-25* mutants and wild-type animals at 4 hours (Fig 2B). Here we found similar levels of IPR gene induction in *pals-22 pals-25* and wild-type animals, suggesting that *pals-22*/*25* regulate expression of IPR genes in parallel to infection. Next, we examined the role of *pals-22*/*25* in the transcriptional response to proteasome blockade, which is another trigger of IPR gene expression [9] (Fig 2C). We used bortezomib, which is a small molecule inhibitor of the 26S proteasome. Here again, we found that bortezomib treatment induced IPR gene expression in *pals-22 pals-25* mutants at levels similar to wild-type animals. Therefore *pals-22/25* appear to regulate IPR gene expression in parallel to infection and proteasomal stress.

### *pals-25* mutations reverse multiple physiological phenotypes caused by *pals-22* mutations

*pals-22* mutants have several striking physiological phenotypes, including slowed growth and shorter lifespans, as well as increased resistance to proteotoxic stress like heat shock [6]. Therefore, we investigated whether mutations in *pals-25* suppress these phenotypes of *pals-22* mutants. First, we investigated developmental rate by measuring the fraction of animals that reach the fourth larval (L4) stage by 48 hours after embryogenesis. Nearly all wild-type animals are L4 at this timepoint, whereas less than 20% of *pals-22* mutants are L4 (Fig 3A). We found that mutations in *pals-25* completely reverse this delayed development of *pals-22* mutants, with nearly all *pals-22 pals-25* mutants reaching the L4 stage by 48 hours (Fig 3A). Next, we analyzed lifespan, as previous work showed that *pals-22* mutants have a significantly shortened lifespan compared to wild-type animals [6, 8]. Here again we found that *pals-25* mutations reversed this effect, with *pals-22 pals-25* mutants having lifespans comparable to wild-type animals (Fig 3B, S3A-B Fig). Next, we investigated the effect of *pals-25* mutations on the thermotolerance capacity of *pals-22* mutants, which is greatly enhanced compared to wild-type animals. We found that *pals-22 pals-25* double mutants have survival after heat shock at levels similar to wild-type animals (Fig 3C, S3C-D Fig), indicating that *pals-25* is required for the enhanced thermotolerance of *pals-22* mutants. Thus, these results show that in a *pals-22* mutant background, wild-type *pals-25* is required to delay development, shorten lifespan and enhance thermotolerance.

**Fig 3.**
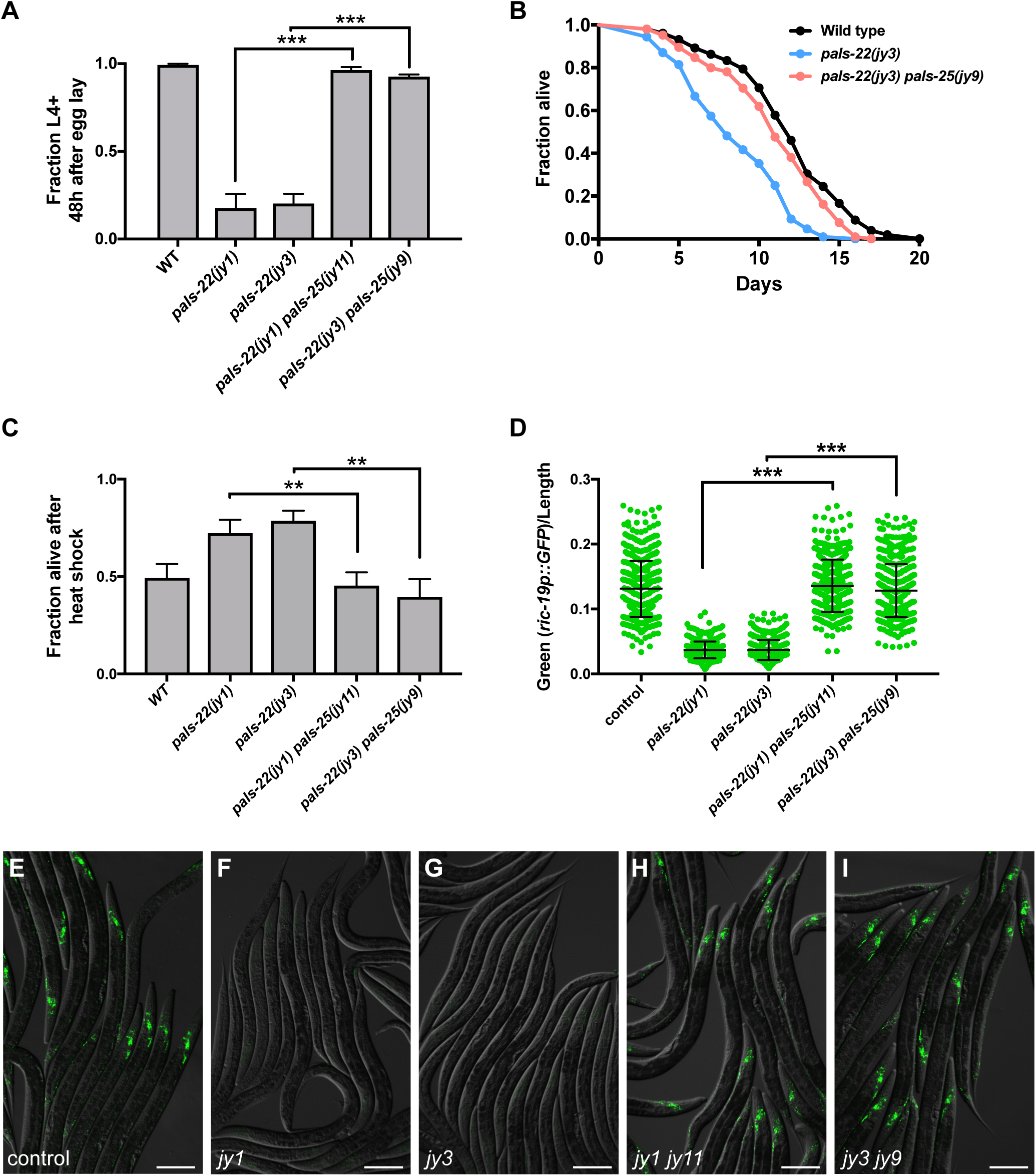
*pals-25* mutations suppress diverse phenotypes of *pals-22* mutants. (A) *pals-25* mutation suppresses the developmental delay of *pals-22* mutants. Fraction of animals reaching the L4 larval stage 48 hours after eggs were laid is indicated. Results shown are the average of three independent biological replicates, with 100 animals assayed in each replicate. Error bars are SD. *** p < 0.001 with Student’s t-test. (B) Lifespan of wild type, *pals-22(jy3)*, and *pals-22(jy3) pals-25(jy9)* animals. Assays were performed with 40 animals per plate, and three plates per strain per experiment. This experiment was repeated three independent times with similar results, with data from a representative experiment shown. See Figures S2A and S2B for other replicates. p-value for *pals-22(jy3)* compared to *pals-22(jy3) pals-25(jy9)* is <0.0001 using the Log-rank test. (C) The increased survival of *pals-22* mutants after heat shock is suppressed by mutations of *pals-25*. Animals were incubated for 2 hours at 37°C followed by 24 hours at 20°C, and then assessed for survival. Strains were tested in triplicate, with at least 30 animals per plate. Mean fraction alive indicates the average survival among the triplicates, errors bars are SD. ** p < 0.01. Assay was repeated three independent times with similar results, and data from a representative experiment are shown. See Figures S2C and S2D for other replicates. (D-I) *pals-25* mutation suppresses transgene silencing in *pals-22* mutants. (D) *ric-19p::GFP* expression quantified in *pals-22* and *pals-22 pals-25* mutants using a COPAS Biosort machine to measure the mean GFP signal and length of individual animals, indicated by green dots. Mean signal of the population is indicated by black bars with error bars as SD. Graph is a compilation of three independent replicates, with at least 100 animals analyzed in each replicate. *** p < 0.001 with Student’s t-test. In (E-I), green is neuronal expression of *ric-19p::GFP*. Shown are (E) wild-type, (F) *pals-22(jy1)*, (G) *pals-22(jy3)*, (H) *pals-22(jy1) pals-25(jy11)*, and (I) *pals-22(jy3) pals-25(jy9)* animals. Images are overlays of green and Nomarski channels and were taken with the same camera exposure for all. Scale bar, 100 *µ*m.

Previous work from the Hobert lab identified *pals-22* in a screen for regulators of reporter gene expression in neurons [8]. They found that mutations in *pals-22* led to decreased levels of GFP reporter expression in neurons and other tissues, and wild-type *pals-22* thus acts as an ‘anti-silencing’ factor of multi-copy transgene expression. Therefore, we analyzed the effects of *pals-25* mutations on transgene silencing in *pals-22* mutants. Here we found that *pals-25* mutations reverse the enhanced silencing of a neuronally expressed GFP transgene in *pals-22* mutants (Fig 3D-I), indicating that wild-type *pals-25* activity is required to silence expression from multi-copy transgenes in a *pals-22* mutant background. Of note, previous work found that a *pals-25* mutation alone does not affect transgene silencing [8]. In summary, mutations in *pals-25* appear to fully reverse all previously described phenotypes of *pals-22* mutants.

### *pals-22* mutants have immunity against coevolved intestinal pathogens of the intestine, which is suppressed by *pals-25* mutations

In addition to the previously described phenotypes of *pals-22* mentioned above, we analyzed resistance of these mutants to intracellular infection. First we analyzed the resistance of *pals-22* mutants to *N. parisii* infection. We fed animals a defined dose of microsporidia spores and measured pathogen load inside intestinal cells. We analyzed pathogen load at 30 hours post infection (hpi), when *N. parisii* is growing intracellularly in the replicative meront stage, and found greatly lowered pathogen load in *pals-22* mutants compared to wild-type animals (Fig 4A-F). We then tested *pals-22 pals-25* double mutants and found these animals to have resistance comparable to wild-type. One explanation for the altered levels of *N. parisii* observed in the intestines of *pals-22* mutant animals is that these mutants have lowered feeding or accumulation of pathogen in the intestine, and thus simply have a lower exposure to *N. parisii*. To address this concern, we added fluorescent beads to our *N. parisii* infection assay and measured accumulation in the intestinal lumen. Here we found that *pals-22* mutants and *pals-22 pals-25* double mutants accumulated fluorescent beads at comparable levels to wild-type animals (S4A Fig), suggesting that their pathogen resistance to *N. parisii* is not simply due to lowered exposure to the pathogen in the intestinal lumen. As a positive control in this assay we tested *eat-2(ad465)* mutants and found that they had reduced fluorescent bead accumulation, consistent with their previously characterized feeding defect [13]. Altogether, these results indicate that *pals-22* and *pals-25* regulate resistance to infection by microsporidia.

**Figure 4.**
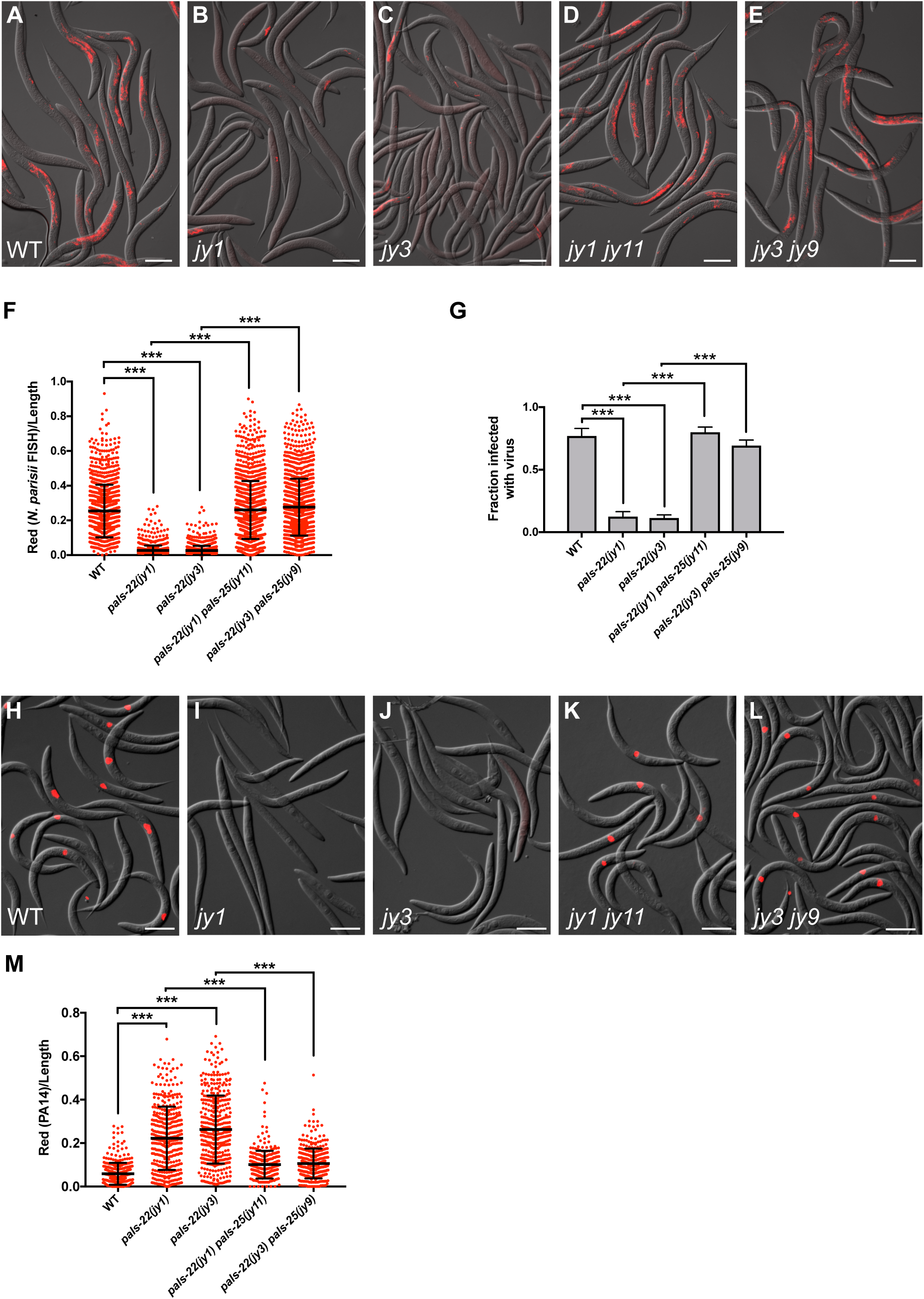
*pals-22* mutants have increased resistance to infection by *N. parisii* or Orsay virus, dependent on *pals-25*. (A-E) Images of (A) wild-type, (B) *pals-22(jy1)*, (C) *pals-22(jy3)*, (D) *pals-22(jy1) pals-25(jy11)*, and (E) *pals-22(jy3) pals-25(jy9)* animals infected with *N. parisii* as L1s, fixed 30 hours post infection, and stained by FISH with an *N. parisii*-specific probe (red). Scale bar, 100 *µ*m. (F) *N. parisii* FISH signal quantified using a COPAS Biosort machine to measure the mean red signal and length of individual animals, indicated by red dots. Mean signal of the population is indicated by black bars, with error bars as SD. Graph is a compilation of three independent replicates, with at least 100 animals analyzed in each replicate. *** p < 0.001 with Student’s t-test. (G) Fraction of animals infected with the Orsay virus 18 hours post infection is indicated. Animals were fixed and stained by FISH with a virus-specific probe, and scored visually for infection. Results shown are the average of three independent biological replicates, with 100 animals assayed in each replicate. Error bars are SD. *** p < 0.001 with Student’s t-test. (H-L) Images of (H) wild-type, (I) *pals-22(jy1)*, (J) *pals-22(jy3)*, (K) *pals-22(jy1) pals-25(jy11)*, and (L) *pals-22(jy3) pals-25(jy9)* animals infected with the Orsay virus as L1s, fixed 18 hours post infection, and stained by FISH with a virus-specific probe (red). Scale bar, 100 *µ*m. (M) Quantification of dsRed fluorescence levels in wild-type, *pals-22*, and *pals-22 pals-25* animals after 16 hours of exposure to dsRed-expressing PA14. Red fluorescence was measured using a COPAS Biosort machine to measure the mean red signal and length of individual animals, indicated by red dots. Mean signal of the population is indicated by black bars, with error bars as SD. Graph is a compilation of three replicates, with at least 100 animals analyzed in each replicate. *** p < 0.001 with Student’s t-test.

We also investigated resistance of *pals-22* mutants and *pals-22 pals-25* double mutants to other pathogens. First, we measured resistance to infection by the Orsay virus. Like *N. parisii*, Orsay virus is a natural pathogen of *C. elegans*, and replicates inside of *C. elegans* intestinal cells [14]. We used FISH staining of Orsay viral RNA to quantify the fraction of worms infected at 18 hpi. Here we found that *pals-22* mutants had significantly decreased viral load when compared to wild-type animals (Fig 4G-L). This lowered viral infection in *pals-22* mutants was fully reversed in *pals-22 pals-25* mutants back to wild-type levels. Importantly, we confirmed that *pals-22* and *pals-22 pals-25* mutants do not have altered fluorescent bead accumulation in the intestine compared to wild-type animals in the presence of virus (S4B Fig), indicating that their lowered viral load is not likely due to lowered exposure to the virus.

Interestingly, we found that *pals-22* mutants did not have reduced pathogen loads when infected with the Gram-negative bacterial pathogen *Pseudomonas aeruginosa* (clinical isolate PA14) (Fig 4M). In fact, these mutants had increased pathogen load, which was suppressed by mutations in *pals-25*. To our knowledge *P. aeruginosa* species are not common pathogens of nematodes in the wild, although under laboratory conditions, *P. aeruginosa* PA14 does accumulate in the *C. elegans* intestinal lumen and causes a lethal infection [15]. In summary *pals-22* mutants have increased resistance to natural pathogens of the intestine, but increased susceptibility to PA14, a ‘non-natural’ pathogen of the intestine.

### RNA-seq analysis of *pals-22/25*-upregulated genes define the IPR

Previous work indicated that *N. parisii* and the Orsay virus induce a common set of genes, despite these being very different pathogens [9]. We called eight of these genes the IPR subset [6], and here we show they are regulated by *pals-22/25* (Fig 2A). To identify additional genes regulated by *pals-22/25*, we performed RNA-seq analysis of *pals-22* mutants, *pals-22 pals-25* mutants, and wild-type animals. We performed differential gene expression analysis using a FDR<0.01 cutoff (see Materials and Methods for a complete description of criteria) and determined that 2,756 genes were upregulated in *pals-22* mutants compared to wild-type animals (Fig 5A, S7 Table). Next we compared *pals-22* mutants to *pals-22 pals-25* double mutants and found that 744 genes were upregulated (Fig 5A, S7 Table). Of these two comparisons, there are 702 genes in common that are upregulated both in *pals-22* mutants compared to wild-type animals and in *pals-22* mutants compared to *pals-22 pals-25* double mutants (Fig 5A). Therefore, these 702 genes are negatively regulated by wild-type *pals-22* and require the activity of the wild-type *pals-25* for their induction in the absence of *pals-22.* These 702 genes include genes like our *pals-5* reporter (Fig 1) and other IPR genes (Fig 2).

**Fig 5.**
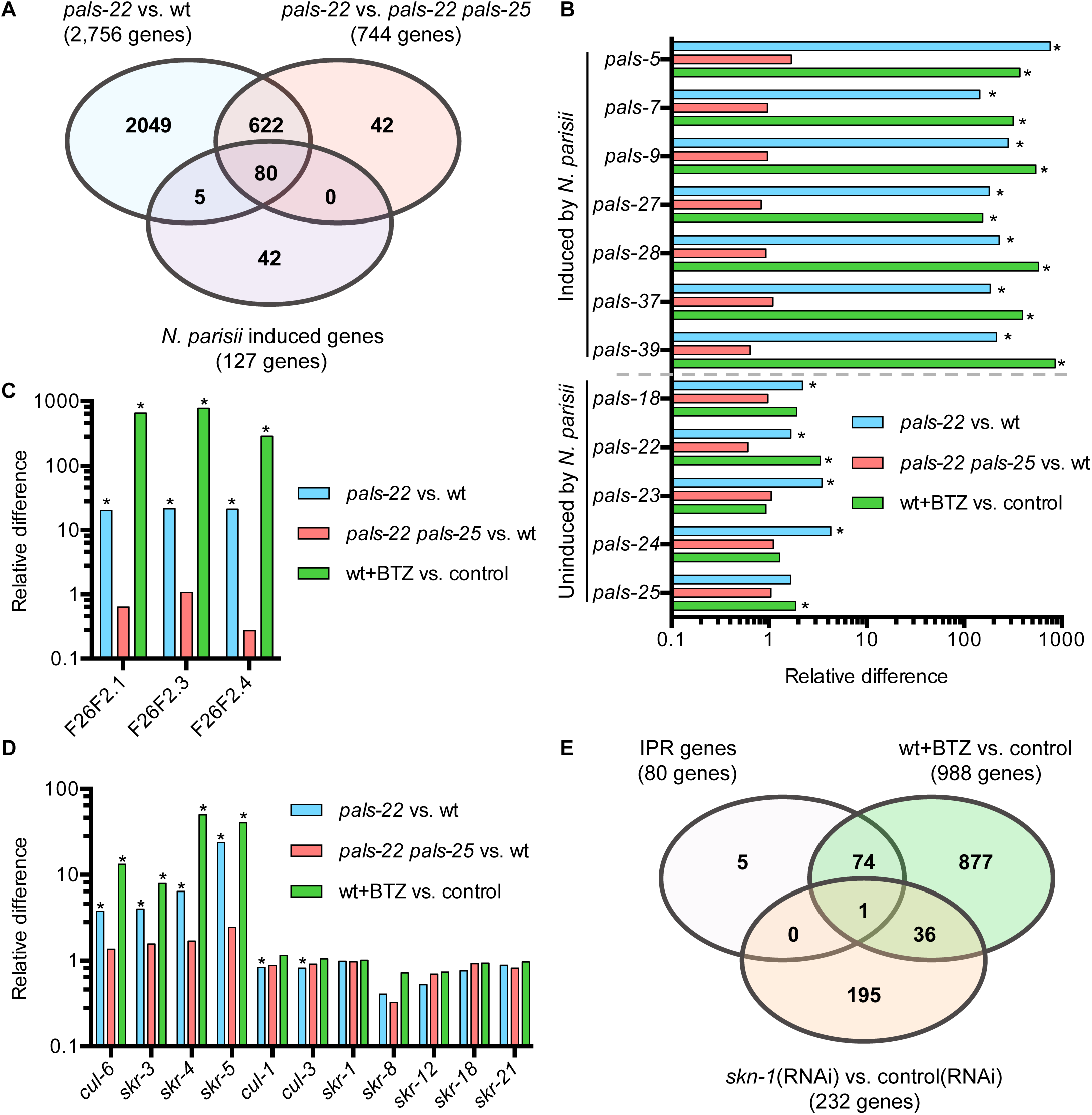
The *pals-22/25* gene pair transcriptionally regulates genes that are induced by *N. parisii* infection or proteasome blockade. (A) Venn diagram comparing 1) genes upregulated in *pals-22* mutants compared to wild-type animals, 2) genes upregulated in *pals-22* mutants compared to *pals-22 pals-25* double mutants and 3) genes induced in wild-type animals in response to *N. parisii* infection. Gene sets 1 and 2 were obtained from RNA-seq data outlined in this study, and Gene set 3 was obtained via RNA-seq in a previous study [9]. We define the IPR genes as the 80 genes common across the three gene sets. (B-D) The relative mRNA levels measured by RNA-seq in: *pals-22(jy3)* animals compared to wild-type; *pals-22(jy3) pals-25(jy9)* animals compared to wild-type; and wild-type animals treated with the proteasome inhibitor bortezomib (BTZ) compared to DMSO (vehicle control for BTZ). * FDR<0.01 as calculated by edgeR and limma (see Materials and Methods) indicates the gene is considered to be differentially expressed. (B) *pals* genes induced by *N. parisii* infection are also regulated by *pals-22/25* and induced by BTZ. (C) Species-specific F26F2 genes are regulated by the *pals-22/25* gene pair and are induced by BTZ. (D) The Cullin-Ring Ligase components Cullin (*cul)* and Skp-related (*skr)* genes that are upregulated during *N. parisii* infection are also regulated by the *pals-22/25* gene pair and induced by BTZ. (E) Venn Diagram showing overlap between: IPR genes defined in (A); genes upregulated by treatment with BTZ; and genes upregulated due to *skn-1* RNAi [16]. See Tables S2 and S3 for detailed expression levels of genes discussed here.

We next compared these 702 *pals-22/25* regulated genes to genes induced during *N. parisii* infection identified in a previous study [9] to expand our list of IPR genes. Out of 127 genes induced during *N. parisii* infection we found that the *pals-22/25* gene pair regulated mRNA expression of 80 of these genes (Fig 5A). Specifically, of the 25 *pals* genes induced upon intracellular infection, all are induced in *pals-22* mutants and reverted back to wild-type levels in *pals-22 pals-25* double mutants (Fig 5B, S7 Table). Notably, all *pals* genes that are not regulated by *pals-22/25* are also not induced by infection. Furthermore, the other nematode-specific genes F26F2.1, F26F2.3, and F26F2.4, which are induced by *N. parisii* and Orsay virus infection, were also found to be induced in *pals-22* mutants and brought back to wild-type levels in *pals-22 pals-25* double mutants (Fig 5C). In addition, we found that the ubiquitin ligase components are similarly regulated (Fig 5D). These studies thus define IPR genes as the 80 genes that are: 1) induced by *N. parisii* infection, 2) induced in a *pals-22* mutant background, and 3) reversed back to wild-type levels in *pals-22 pals-25* double mutants.

### Genes regulated by *pals-22/25* and infection are also regulated by proteasomal stress

Previous work indicated that blockade of the proteasome, either pharmacologically or genetically, will induce expression of a subset of IPR genes [9]. To determine the IPR genes that are induced by proteasome stress, we performed RNA-seq analysis to define the whole-genome response to this stress. Again, we conducted differential expression analysis and compared gene expression of animals after 4 hours of exposure to the proteasome inhibitor bortezomib compared to the DMSO vehicle control. From these experiments we determined that 988 genes are induced following bortezomib treatment, using the cut-off mentioned above and described in the Materials and Methods. Interestingly, nearly all of the IPR genes described above are also induced following bortezomib treatment (Fig 5E). Previous work has shown that genes induced by *N. parisii* do not include the proteasome subunits induced by proteasome blockade as part of the bounceback response [9]. The bounceback response is induced via the transcription factor SKN-1. Consistent with these results, here we find that the IPR genes induced by bortezomib are distinct from those regulated by the transcription factor SKN-1, as defined by a previous study [16]. The overlap between SKN-1 regulated genes and IPR genes includes only one gene (Fig 5E).

As shown earlier, *pals-22* mutants have increased resistance to heat shock, and previous work indicated that there is overlap between genes induced by chronic heat stress and genes induced by *N. parisii* and virus infection [9]. However, the genes in common are distinct from the canonical chaperones, or Heat Shock Proteins (HSPs), which are induced by the heat shock transcription factor HSF-1. To learn more about the connection between heat shock response, HSF-1, and the IPR, we compared the IPR genes with those induced by HSF-1 as defined in a previous study [17]. Here we found 8 genes in common between our set of 80 IPR genes and the set of 368 genes upregulated by HSF-1, none of which are predicted to encode chaperone proteins (S10 Table). We also compared the 368 genes upregulated by HSF-1 with the 702 genes that are regulated by *pals-22/25* and found 59 genes in common (S10 Table). These genes include secreted C-type lectins and F-box genes, but do not include chaperones. In summary, HSF-1 regulates 59 genes in common with those regulated by *pals-22/25*, but only 8 of these are IPR genes.

### *pals-22* and *pals-25* regulate expression of genes induced by other natural pathogens

Next, we used Gene Set Enrichment Analysis (GSEA) to broadly compare *pals-22/25*-regulated genes to genes regulated by other pathogens, stressors, and stress-related pathways. Here we found that *pals-22/25* does not significantly regulate expression of genes induced by the Gram-negative bacterial pathogen *P. aeruginosa* or the Gram-positive bacterial pathogens *Staphylococcus aureus* and *Enterococcus faecalis* as analyzed in previous studies (Fig 6). Notably, the strains used in these studies are clinical isolates. Furthermore, these pathogen species are not known to be natural pathogens of nematodes and are not found inside *C. elegans* intestinal cells before there is extensive tissue damage in the host [18]. (Refer to S9 Table for additional comparisons among genes regulated by *pals-22/25*, bortezomib treatment, and other pathogens and stress pathways.)

**Fig 6.**
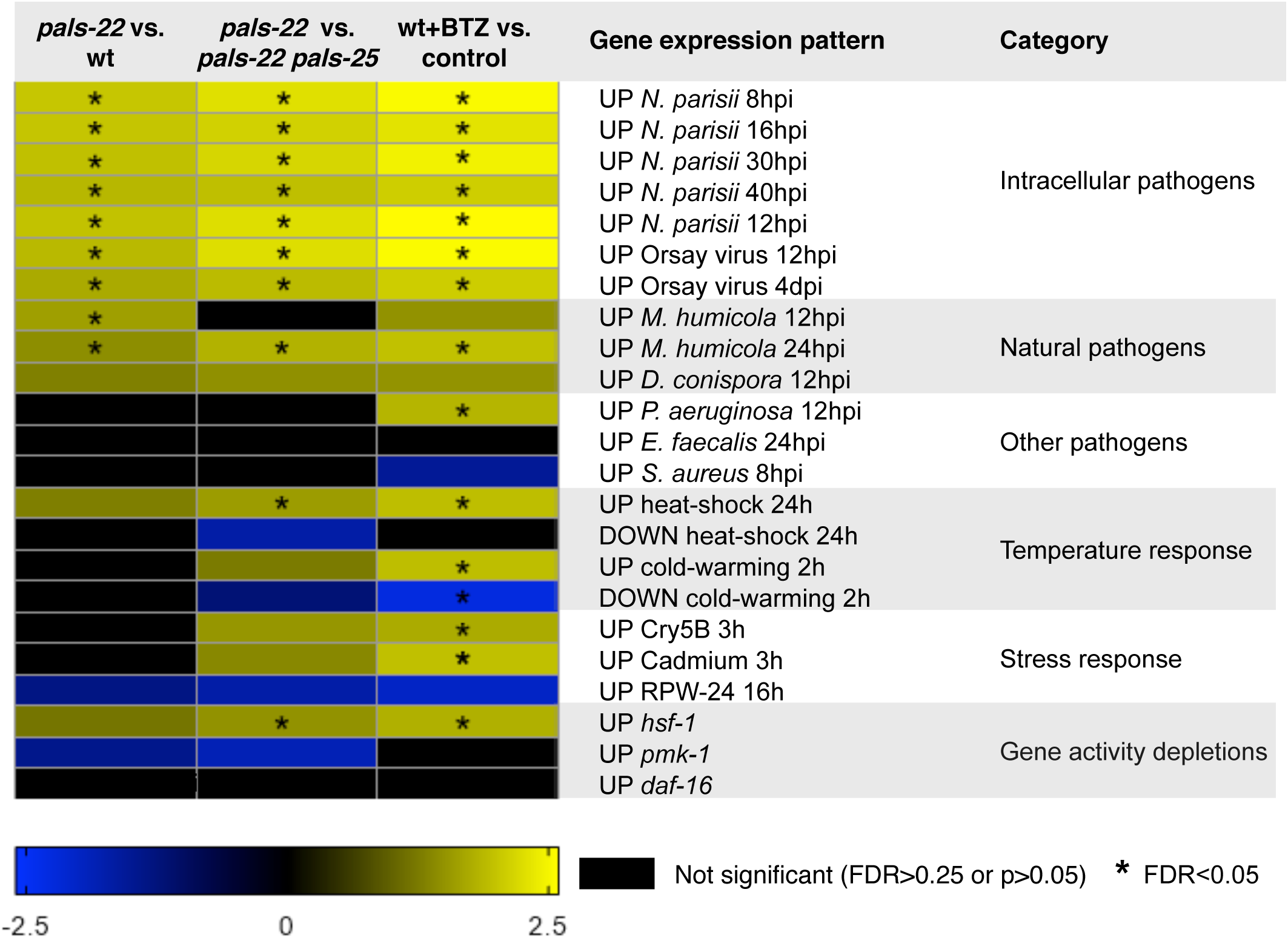
Functional analysis of genes transcriptionally regulated by the *pals-22/25* gene pair. Correlation of genes transcriptionally regulated by the *pals-22/25* gene pair and genes differentially expressed due to bortezomib treatment with genes that are up-or downregulated in response to infection by pathogens or other environmental stresses. Analysis was performed using the GSEA 3.0 software package (see Materials and Methods) and correlation is quantified as a Normalized Enrichment Score (NES). A positive NES (yellow) indicates correlation with upregulated genes in the denoted comparison while a negative NES (blue) indicates correlation with downregulated genes. Black cells indicate no significant correlation was detected, a FDR>0.25, or p>0.05). *FDR<0.05. For more detailed results, see S9 Table. For details on the gene sets used see S8 Table.

Because *pals-22* and *pals-25* regulate expression of a majority of the genes induced by natural intestinal pathogens like *N. parisii* and the Orsay virus, we investigated whether they regulate the transcriptional response to natural pathogens that infect other tissues. The fungal pathogen *Drechmeria coniospora* infects and penetrates the epidermis of nematodes, triggering a GPCR signaling pathway that upregulates expression of neuropeptide-like (*nlp*) genes including *nlp-29* to promote defense [15]. Our transcriptome analysis shows that *pals-22/25* regulate a significant number of genes in common with *Drechmeria* infection (S10 Table). Notably these genes do not include the well-characterized neuropeptide *nlp* defense genes, although they do include many of the *pals* genes. A more recently described natural pathogen of *C. elegans* is *Myzocytiopsis humicola*, which is an oomycete that also infects through the epidermis and causes a lethal infection [19]. Here as well, *pals-22/25* regulate a significant number of genes in common with those induced by *M. humicola* infection, including the <u>chi</u>tinase-like ‘*chil’* genes that promote defense against this pathogen (S10 Table). Interestingly, these *chil* genes, like the *pals* genes, are species-specific [8, 19].

We next used Ortholist [20] to determine which genes identified from our RNA-seq analyses have predicted human orthologs. Of the 702 genes regulated by *pals-22/25*, 279 genes (39.7%) have predicted human orthologs (S11 Table). In contrast, of the 368 genes induced in *hsf-1* mutants 190 (51.6%) have predicted human orthologs. Therefore, more of the genes regulated by the conserved transcription factor *hsf-1* have human orthologs compared to genes regulated by the *C. elegans*-specific *pals-22/25* gene pair. Furthermore, when we restrict our analysis to just the 80 IPR genes, only 14 (17.5%) have predicted human orthologs, indicating that the transcriptional response to natural infection is enriched for genes that are not well-conserved.

### *pals-22/25* control expression of epidermal defense genes induced by oomycetes

As described above, the RNA-seq analysis of genes regulated by *pals-22/25* indicated that this gene pair controls expression of genes induced by diverse natural pathogens of *C. elegans*. Indeed, in a forward genetic screen for *C. elegans* genes that regulate expression of the *M. humicola*-induced *chil-27p::GFP* reporter, we isolated independent loss-of-function alleles of *pals-22* (Fig 7A). These mutant alleles cause constitutive expression of *chil-27p::GFP* in the epidermis in the absence of infection (Fig 7B). RNAi against *pals-22* also led to constitutive GFP expression (S12 Fig), in a manner that is indistinguishable from that observed upon infection with *M. humicola*. These results indicate that *pals-22* acts as a negative regulator of *chil-27* expression in the epidermis.

**Fig 7.**
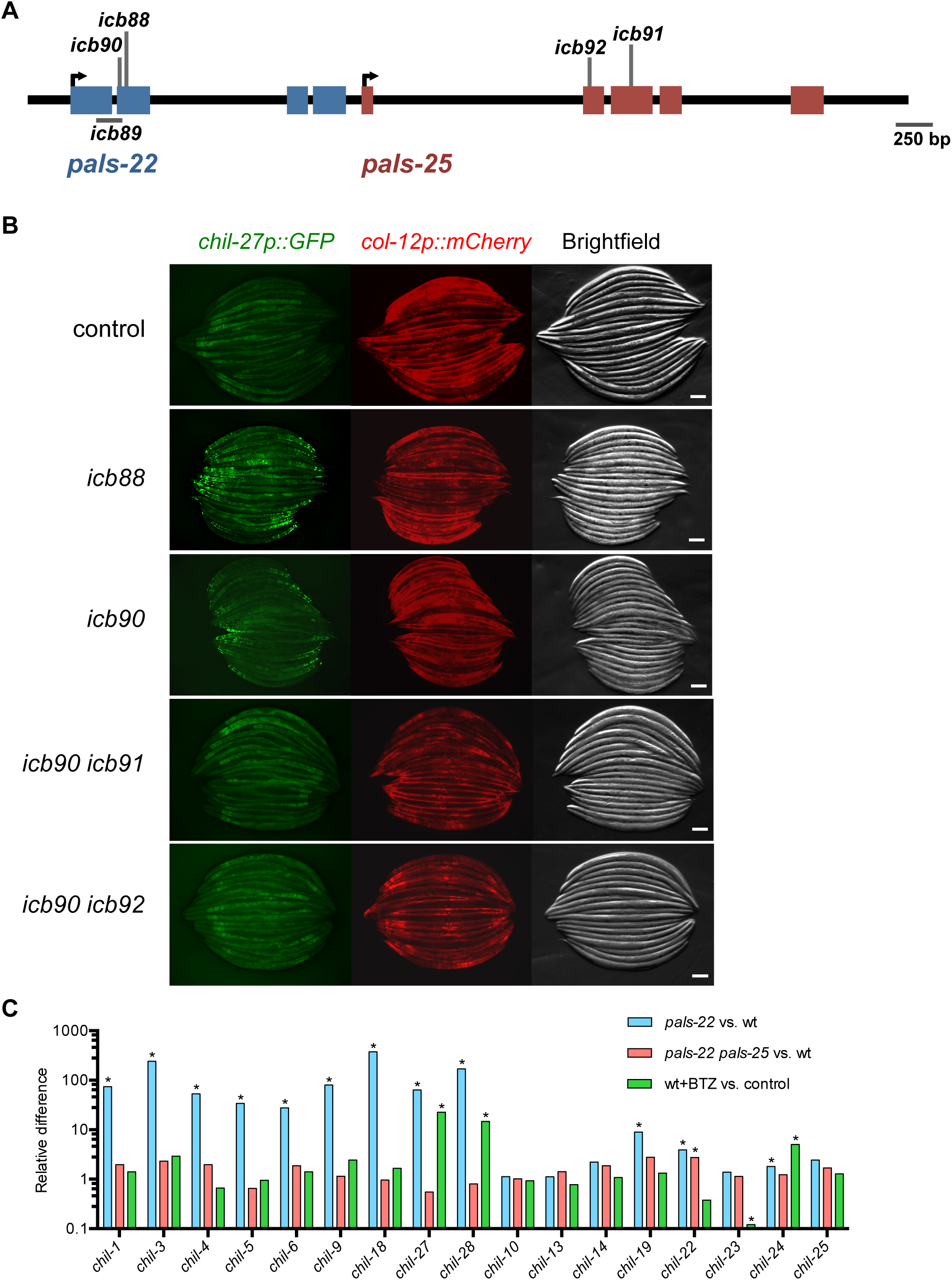
*pals-22* and *pals-25* regulate expression of *chil-27* in the epidermis. (A) *pals-22* and *pals-25* gene coding structure (UTR not shown), with blue exons for *pals-22* and red exons for *pals-25*. See S1 Table for residues altered. (B) Expression of *chil-27p::GFP* is regulated by *pals-22* and *pals-25*. Shown are control, *pals-22(icb88), pals-22(icb90), pals-22(icb90) pals-25(icb91)*, and *pals-22(icb90) pals-25(icb92)* animals. The *col-12p::mCherrry* transgene is constitutively expressed in the epidermis. Scale bar, 100 *µ*m. (C) Relative levels of *chil* gene expression. *pals-22/25* regulate *chil* genes that are induced upon infection by oomycete. *FDR<0.01 as calculated by edgeR and limma (see Materials and Methods) indicates the gene is considered to be differentially expressed.

We then used a *pals-22; chil-27p::GFP* strain for a suppressor screen, analogous to the one described earlier for suppressors of GFP expression in *pals-22; pals-5p::GFP* (Fig 1). Interestingly, in this new screen we isolated two new alleles of *pals-25* which fully suppress the constitutive gene expression of *chil-27p::GFP* seen in *pals-22* mutants (Fig 7A-B), indicating that wild-type *pals-25* acts as a positive regulator of *chil-27* expression. These observations are consistent with our differential expression analysis, which determined that *chil-27* is induced in a *pals-22* mutant background and that *pals-25* is required for this induction (Fig 7C). Therefore, *pals-22/25* act as a switch not only for genes induced in the intestine by natural intestinal pathogens, but also as a switch for genes induced in the epidermis by natural epidermal pathogens of *C. elegans*.

## Discussion

In many organisms, there is a balance between growth and pathogen resistance. In particular, many studies in plants have indicated that genetic immunity to disease comes at a cost to the yield of crops [21]. Here we define a program in *C. elegans* that controls a balance between organismal growth with resistance to natural pathogens, which is regulated by the *pals-22/25* species-specific gene pair. These genes act as a switch between a ‘defense program’ of enhanced resistance against diverse natural pathogens like microsporidia and virus, improved tolerance of proteotoxic stress and increased defense against exogenous RNA [8], and a ‘growth program’ of normal development and lifespan (Fig 8). We call this physiological defense program the “IPR” and it appears to be distinct from other canonical stress response pathways in *C. elegans*, including the p38 MAP kinase pathway, the insulin-signaling pathway, and the heat shock response, among others [6]. Our previous analyses indicated that ubiquitin ligases may play a role in executing the IPR program, as the cullin/CUL-6 ubiquitin ligase subunit is required for the enhanced proteostasis capacity of *pals-22* mutants [6].

**Fig 8.**
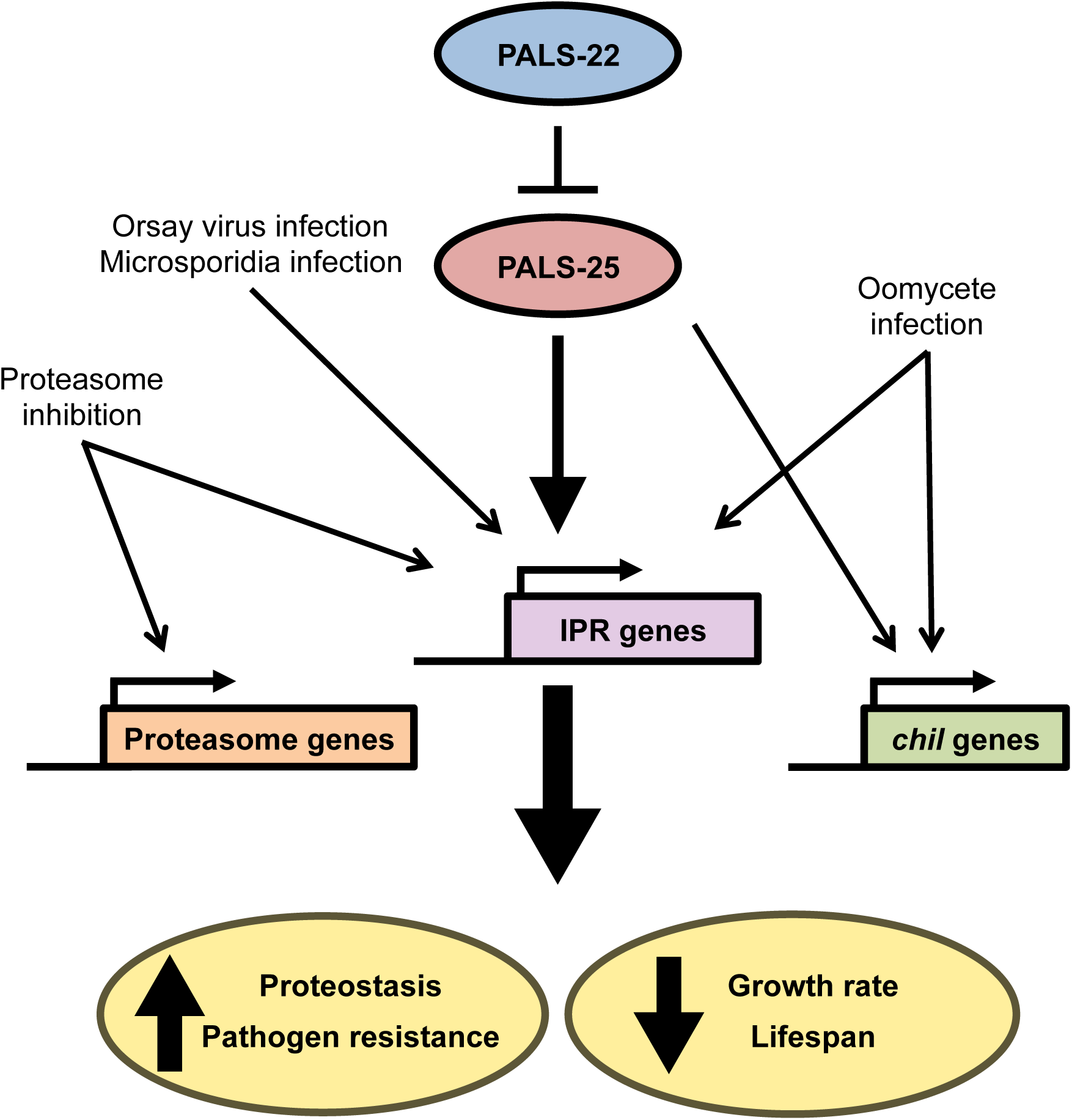
Model for *pals-22/pals-25* regulation of response to natural pathogens of *C. elegans*.

*pals-22* mutants are highly resistant to the microsporidian pathogen *N. parisii*, which is the most common parasite found in wild-caught *C. elegans* [22, 23]. Little is known about innate immune pathways that provide defense against *N. parisii*. Canonical immune pathways in *C. elegans* like the p38 MAP kinase pathway provide defense against most pathogens tested in *C. elegans* but do not provide defense against *N. parisii* [12]. The mechanism by which *pals-22/25* regulate resistance to *N. parisii* is not clear. Our RNA-seq analysis demonstrates that *pals-22/25* affect expression of hundreds of genes in the genome. In particular, most of the genes induced by the natural intracellular pathogens Orsay virus and *N. parisii* are controlled by *pals-22* and *pals-25*, although the function of these IPR genes in defense is unknown. Interestingly, we found that *pals-22/25* regulate expression of genes induced not only by natural intestinal pathogens but also of genes induced by natural epidermal pathogens, such as the oomycete species *M. humicola. M. humicola* induces expression of *chil-*gene family, and genetic analysis shows these genes promote defense against *M. humicola* [19]. Notably, we identified *pals-22/25* in independent forward genetic screens for regulators of *chil-27* and found that they regulate expression of this defense gene in the epidermis. Thus, *pals-22/25* regulate expression of genes induced by diverse natural pathogens.

While *pals-25* is required to activate IPR gene expression in a *pals-22* mutant background, it is not required for activation of IPR gene expression in response to *N. parisii* infection or proteasomal stress. Therefore, *pals-22/25* may not mediate detection of these pathogens, although they might mediate detection and be redundant with other factors. Intriguingly, the *pals-22/25* gene pair share evolutionary and phenotypic features with plant R gene pairs, which serve as sensors for virulence factors delivered into host cells by co-evolved plant pathogens. For example, the *Arabidopsis thaliana* gene pair RRS1 and RPS4 are species-specific, share the same promoter, and direct opposite outcomes, with RRS1 inhibiting and RPS4 promoting ‘effector-triggered immunity’ against natural pathogens [24, 25]. Similarly, *pals-22* and *pals-25* are species-specific, appear to be in an operon together, and direct opposite physiological outcomes including defense against natural pathogens. RRS1 and RPS4 proteins directly bind to each other, and RRS1 normally inhibit RPS4 function until detection of bacterial virulence factors, at which point RRS1 inhibition is relieved and RPS4 is free to promote pathogen defense, although the steps downstream of RRS1/RPS4 are poorly understood. Although the *pals* genes do not share sequence similarity with the R genes, in this analogy PALS-22 would inhibit PALS-25 and serve as the ‘tripwire’ to detect virulence factors from natural pathogens and free PALS-25 to promote the IPR defense program. While this model is attractive, it is purely speculative as we currently have no direct evidence that PALS-22 detects virulence factors. Identification of such hypothetical virulence factors would be the focus of future studies.

The molecular events by which *C. elegans* detects infection are poorly understood, although nematodes do appear to use a form of effector-triggered immunity or ‘surveillance immunity’. Studies with several distinct pathogens have indicated that *C. elegans* induces defense gene expression in response to perturbation of core processes like translation and the ubiquitin-proteasome system [26]. For example, studies with *P. aeruginosa* demonstrated that *C. elegans* detects the presence of the translation-blocking Exotoxin A through its effects on host translation, not through detection of the shape of the toxin [27, 28]. In addition to this mode of detection, *C. elegans* may also detect specific molecular signatures like canonical Pathogen-Associated Molecular Patterns (PAMPs). In all likelihood, several types of pathogen detection are used by *C. elegans*. Surprisingly however, there have been no direct PAMP ligand/receptor interactions demonstrated for pattern recognition receptors (PRR) in the worm, although there has been a Damage-Associated Molecular Pattern (DAMP)/G-protein-coupled receptor interaction demonstrated to be critical for response to *Drechmeria* infection [29]. Indeed, *C. elegans* lacks many PRR signaling pathways that are well described in flies and mammals. For example, the *C. elegans* single Toll-like receptor *tol-1* does not act canonically and worms appear to have lost its downstream transcription factor NFkB, which is critical for innate immunity in flies and mammals [30]. Perhaps conservation of immune genes is only reserved for defense against rare, ‘non-natural’ pathogens, because genes that are important for immunity are subject to attack and inhibition by microbes [31]. Thus, immune genes that provide defense against natural pathogens from the recent evolutionary past will not be broadly conserved but rather will be species-specific, like rapidly evolving R genes in plants. While R genes have been shown to encode proteins that detect virulence factors secreted into host cells by co-evolved plant pathogens, the mechanism by which they activate downstream immune signaling is unclear. We propose that the IPR physiological program regulated by the *pals-22/25* antagonistic paralogs in *C. elegans* could be analogous to effector-triggered immunity regulated by opposing R gene pairs like RRS1/RPS4 used in plants for resistance against co-evolved pathogens.

Interestingly, an example of vertebrate-specific antagonistic paralogs has recently been described to play a role in regulating nonsense-mediated RNA decay (NMD) [32]. These studies provide a potential explanation to the long-standing question of how gene duplications are retained, when they are presumably redundant immediately following gene duplication. Specifically, this model predicts that gene duplication events can be rapidly retained if the proteins made from these genes are involved in protein-protein interactions. With just one non-synonymous nucleotide change that switches a wild-type copy to become dominant negative within a multimeric signaling complex, a gene duplication event can be selected for and retained in the heterozygote state – i.e. in one generation. Perhaps in this way, new genes can be born and survive, when gene pairs can evolve to direct opposing functions like the Upf3a/3b paralogs in NMD, and the RRS1/RPS4 and *pals-22/25* paralogs in immunity/growth.

## Methods

### Strains

*C. elegans* were maintained at 20°C on Nematode Growth Media (NGM) plates seeded with Streptomycin-resistant *E. coli* OP50-1 bacteria according to standard methods [33]. We used N2 wild-type animals. Mutant or transgenic strains were backcrossed at least three times. See S1 Table for a list of all strains used in this study.

### EMS screens and cloning of alleles

*pals-22* mutant worms (either the *jy1* or *jy3* allele) carrying the *jyIs8[pals-5p::GFP, myo-2p::mCherry]* transgene were mutagenized with ethyl methane sulfonate (EMS) (Sigma) using standard procedures as described [34]. L4 stage P0 worms were incubated in 47 mM EMS for 4 hours at 20°C. Worms were screened in the F2 generation for decreased expression of GFP using the COPAS Biosort machine (Union Biometrica).

Complementation tests were carried out by generating worms heterozygous for two mutant alleles and scoring *pals-5p::GFP* fluorescence. For whole-genome sequencing analysis of mutants, genomic DNA was prepared using a Puregene Core kit (Qiagen) and 20X sequencing coverage was obtained. We identified only one gene (*pals-25*) on LGIII containing variants predicted to alter function in both mutants sequenced (*jy9* and *jy100*). Additional *pals-25* alleles were identified by Sanger sequencing. Screens in the strains carrying the *icbIs4[chil-27p::GFP, col-12p::mCherry]* transgene were performed in a similar manner except that we used 24 mM EMS to recover the *pals-22* alleles (*icb88, icb90*) and 17 mM EMS for the *pals-22(icb90)* suppressor screen and that in both cases we selected F2 animals manually using a Zeiss Axio ZoomV16 dissecting scope. The two *pals-22* alleles (*icb88, icb90*) were identified by whole genome sequencing of GFP positive F2 recombinants after crossing to the polymorphic isolate CB4856 as previously described [35] whereas the two *pals-25* alleles were found by direct sequencing of the mutant strains. The *pals-22(icb89)* allele was identified by Sanger sequencing. See S1 Table for a list of all mutations identified.

### RNA interference

RNA interference was performed using the feeding method. Overnight cultures of RNAi clones in the HT115 bacterial strain were seeded onto NGM plates supplemented with 5 mM IPTG and 1 mM carbenicillin and incubated at 25°C for 1 day. Eggs from bleached parents or synchronized L1 stage animals were fed RNAi until the L4 stage at 20°C. For all RNAi experiments an *unc-22* clone leading to twitching animals was used as a positive control to test the efficacy of the RNAi plates. The *pals-22* RNAi clone (from the Ahringer RNAi library) was verified by sequencing. The *pals-25* RNAi clone was made with PCR and includes 1079 base pairs spanning the second, third, and fourth exons of *pals-25*. This sequence was amplified from N2 genomic DNA, cloned into the L4440 RNAi vector, and then transformed into HT115 bacteria for feeding RNAi experiments.

### Quantitative RT-PCR

Endogenous mRNA expression changes were measured with qRT-PCR as previously described [6]. Synchronized L1 worms were grown on NGM plates at 20°C to the L4 stage and then collected in TriReagent (Molecular Research Center, Inc.) for RNA extraction. For *N. parisii* infection, 7 × 10^6^ spores were added to plates with L4 stage worms and then incubated at 25°C for 4 hours before RNA isolation. Bortezomib (or an equivalent amount of DMSO) was added to L4 stage worms for a final concentration of 20 μM; plates were then incubated at 20°C for 4 hours before RNA isolation. At least two independent biological replicates were measured for each condition, and each biological replicate was measured in duplicate and normalized to the *snb-1* control gene, which did not change upon conditions tested. The Pffafl method was used for quantifying data [36].

### Heat shock assay

Worms were grown on standard NGM plates until the L4 stage at 20°C and then shifted to 37°C for two hours. Following heat shock, plates were laid in a single layer on the bench top for 30 minutes to recover, and then moved to a 20°C incubator overnight. Worms were scored in a blinded manner for survival 24 hours after heat shock; animals not pumping or responding to touch were scored as dead. Three plates were assayed for each strain in each replicate, with at least 30 worms per plate, and three independent assays were performed.

### GFP fluorescence measurement

Synchronized L1 stage animals were grown at 20°C to the L4 stage. The COPAS Biosort machine (Union Biometrica) was used to measure the time of flight (as a measure of length) and fluorescence of individual worms. At least 100 worms were measured for each strain, and all experiments were repeated three times.

### Lifespan

L4 stage worms were transferred to 6 cm NGM plates seeded with OP50-1 bacteria and incubated at 25°C. Worms were scored every day, and animals that did not respond to touch were scored as dead. Animals that died from internal hatching or crawled off the plate were censored. Worms were transferred to new plates every day throughout the reproductive period. Three plates were assayed for each strain in each replicate, with 40 worms per plate.

### Microscopy

Worms were anesthetized with 10 μM levamisole in M9 buffer and mounted on 2% agarose pads for imaging. Images in Figure S1B and S1C were captured with a Zeiss LSM700 confocal microscope. All other *C. elegans* images were captured with a Zeiss AxioImager M1 or Axio Zoom.V16.

### N. parisii and Orsay virus infection assays

*N. parisii* spores were prepared as previously described [37], and Orsay virions were prepared as described previously [9]. For pathogen load analysis, synchronized L1 worms were plated with a mixture of OP50 bacteria and 5 × 10^5^ *N. parisii* spores or a 1:20 dilution of Orsay virus filtrate, and then incubated at 25°C for either 30 hours (*N. parisii*) or 18 hours (Orsay virus) before fixing with paraformaldehyde. Fixed worms were stained with individual FISH probes conjugated to the red Cal Fluor 610 dye (Biosearch Technologies) targeting either *N. parisii* ribosomal RNA or Orsay virus RNA. *N. parisii* pathogen load was measured with the COPAS Biosort machine (Union Biometrica). Orsay virus infection was assayed visually using the 10x objective on a Zeiss AxioImager M1 microscope. In feeding measurement assays, plates were set up as for pathogen infection with the addition of fluorescent beads (Fluoresbrite Polychromatic Red Microspheres, Polysciences Inc.). Worms were fixed in paraformaldehyde after 30 minutes and red fluorescence signal was measured with the COPAS Biosort machine (Union Biometrica).

### P. aeruginosa pathogen load

Overnight cultures of a *P. aeruginosa* PA14-dsRed strain [38] were seeded onto SK plates with 50 μg/ml ampicillin, and then incubated at 37°C for 24 hours followed by 25°C for 24 hours. Worms at the L4 stage were washed onto the PA14-dsRed plates, incubated at 25°C for 16 hours, and then assayed with a COPAS Biosort machine (Union Biometrica) for the amount of red fluorescence inside each animal.

### RNA-seq sample preparation

Synchronized L1 stage worms were grown on 10 cm NGM plates seeded with OP50-1 *E. coli* at 20°C until worms had reached the L4 stage. N2, *pals-22(jy3)*, and *pals-22(jy3) pals-25(jy9)* strains were then shifted to 25°C for 4 hours before harvesting for RNA extraction. Bortezomib (or an equivalent amount of DMSO) was added to plates with L4 stage N2 worms for a final concentation of 20uM; plates were then incubated at 20°C for 4 hours before RNA isolation. RNA was isolated with TriReagent purification, followed by RNeasy column cleanup (Qiagen), as described [39]. RNA quality was assessed by Tapestation analysis at the Institute for Genomic Medicine (IGM) at UC San Diego. Paired-end sequencing libraries were then constructed with the TruSeq Stranded mRNA method (Illumina), followed by sequencing on HiSeq4000 machine (Illumina).

### RNA-seq analysis

Sequencing reads were aligned to WormBase release WS235 using Bowtie 2 [40], and transcript abundance was estimated using RSEM [41]. Differential expression analysis was performed in RStudio (v1.1.453) [42] using R (v3.50) [43] and Bioconductor (v3.7) [44] packages. As outlined in the RNAseq123 vignette [45], data was imported, filtered and normalized using edgeR [46], and linear modeling and differential expression analysis was performed using limma [47]. An FDR [48] cutoff of <0.01 was used to define differentially expressed genes; no fold-change criteria was used. Lists of upregulated genes used for comparisons were exported and further sanitized to remove dead genes and update WBGeneIDs to Wormbase release WS263.

### Functional analysis

Functional analysis was performed using Gene Set Enrichment Analysis (GSEA) v3.0 software [49, 50]. Normalized RNA-seq expression data were converted into a GSEA-compatible filetype and ranked using the signal-to-noise metric with 1,000 permutations. Gene sets from other studies were converted to WBGeneIDs according to WormBase release WS263. Independent analyses were performed for each of three comparisons: untreated *pals-22(jy3)* versus untreated N2 animals; untreated *pals-22(jy3)* versus untreated *pals-22(jy3) pals-25(jy9)* animals; bortezomib treated N2 versus DMSO vehicle control treated N2. Results were graphed based on their NES-value using GraphPad Prism 7 (GraphPad Software, La Jolla, CA).

## Acknowledgments

We acknowledge the IGM for RNAseq library construction and sequencing, Corrina Elder, Jessica Sowa, Ivana Sfarcic, and Michael K. Fasseas for technical support, and Vladimir Lazetic, Robert Luallen, Johan Panek, Ivana Sfarcic, Eillen Tecle, Samira Yitiz and Elina Zuniga for comments on the manuscript.

## Supporting information

**S1 Table. Lists of strains and mutations**.

**S2 Figure. *pals-25* RNAi suppresses increased *pals-5p::GFP* expression in *pals-22* mutants and PALS-25::GFP is expressed broadly**.

(A) Wild-type or *pals-22* mutant animals carrying the *pals-5p::GFP* transgene, treated with either L4440 RNAi control or *pals-25* RNAi. Green is *pals-5p::GFP*, red is *myo-2p::mCherry* expression in the pharynx as a marker for presence of the transgene. Images are overlays of green, red, and Nomarski channels and were taken with the same camera exposure for all. Scale bar, 100 *µ*m. (B,C) Confocal fluorescence images of adult animals carrying a fosmid transgene expressing PALS-25::GFP from the endogenous promoter. Animals were treated with either (B) L4440 RNAi control or (C) *pals-22* RNAi. Scale bar, 50 *µ*m.

**S3 Figure. *pals-25* mutation suppresses the lifespan and thermotolerance phenotypes of *pals-22* mutants.**

(A,B) Lifespan of wild type, *pals-22(jy3)*, and *pals-22(jy3) pals-25(jy9)* animals. Assays were performed with 40 animals per plate, and three plates per strain per experiment. p-value for *pals-22(jy3)* compared to *pals-22(jy3) pals-25(jy9)* is <0.0001 using the Log-rank test. (C,D) Survival of animals after 2 hour heat shock treatment at 37°C followed by 24 hours at 20°C. Strains were tested in triplicate, with at least 30 animals per plate. Mean fraction alive indicates the average survival among the triplicates, errors bars are SD. ** p < 0.01, * p < 0.05.

**S4 Figure. Mutation of *pals-22* or *pals-25* does not affect feeding rates of animals in pathogen infection assays.**

(A,B) Quantification of fluorescent bead accumulation in wild-type, *pals-22, pals-22 pals-25*, and *eat-2* mutant animals. Beads were mixed with OP50-1 bacteria and either (A) *N. parisii* spores or (B) Orsay virus and fed to worms as in infection assays. Worms were fixed in paraformaldehyde after 30 minutes of feeding, and fluorescence of accumulated beads in each animal was measured using a COPAS Biosort machine to measure the mean red signal and length of individual animals, indicated by red dots. Mean signal of the population is indicated by black bars, with error bars as SD. Graph is a compilation of three independent replicates, with at least 100 animals analyzed in each replicate. Statistical analysis was performed using one-way ANOVA. *** p < 0.001, ns, not significant.

**S5 Table. RNA-seq statistics.**

**S6 Table. FPKM values for all genes in data set.**

**S7 Table. Differentially expressed genes, as determined by edgeR and limma.**

**S8 Table. Gene sets used for GSEA and their sources.**

**S9 Table. Detailed GSEA results.**

**S10 Table. Gene set overlaps.**

**S11 Table. Human orthology analysis.**

**S12 Figure. Induction of *chil-27p::GFP* expression seen after *pals-22* RNAi treatment.**

Shown are animals treated with either L4440 RNAi control, *pals-22* RNAi, or *M. humicola* infection. The *col-12p::mCherrry* transgene is constitutively expressed in the epidermis. Scale bar, 100 *µ*m.

